# Hypoxia inducible factor-2α increases sensitivity of colon cancer cells towards oxidative cell death

**DOI:** 10.1101/823997

**Authors:** Rashi Singhal, Sreedhar R Mitta, Kenneth P. Olive, Costas A. Lyssiotis, Yatrik M. Shah

**Affiliations:** Department of Molecular and Integrative Physiology, University of Michigan, Ann Arbor, MI 48109; Department of Internal Medicine, Division of Gastroenterology, University of Michigan, Ann Arbor, MI 48109; Rogel Cancer Center, University of Michigan, Ann Arbor, MI 48109; Department of Pathology, Columbia University Medical Center, New York, NY 10032; Division of Digestive and Liver Diseases, Department of Medicine, Columbia University Medical Center, New York, NY 10032; Herbert Irving Comprehensive Cancer Center, Columbia University Medical Center, New York, NY 10032

**Author notes:** **Correspondence:** Yatrik M. Shah, Department of Molecular & Integrative Physiology, Department of Internal medicine, Division of Gastroenterology, University of Michigan Medical School, Ann Arbor, MI 48109.

## Abstract

Colorectal cancer (CRC) is the second leading cause of cancer-related deaths in the US. Hypoxia is a hallmark of solid tumors which promotes tumor cell growth, survival, metastasis and confers resistance to chemo and radiotherapies. Targeting hypoxic cells has been difficult. Moreover, inhibitors for the major transcription factors, hypoxia inducible factor (HIF)-1α and HIF-2α have not shown long-term efficacy in most cancers. We have previously shown that HIF-2α is essential for colon tumorigenesis. Using an unbiased screen, we show a significant increase in synthetic lethality of HIF-2α overexpressing tumor enteroids to oxidative cell death activators. The treatment with hypoxia mimetic FG4592 (Roxadustat), led to a robust increase in erastin-, RSL3-, and dimethyl fumarate-induced cell death in a dose- and time-dependent manner. Further, our in-vitro data shows that HIF-2α knock-down cells are completely resistant to these drugs. HIF activation promotes upregulation of lipid synthesis genes in vitro and in vivo leading to oxidative stress. Taken together, our results suggest that this intrinsic sensitivity towards oxidative stress associated with hypoxia could be utilized as a persistent and dynamic form of cell death for colon cancer treatment.

## Introduction

Colorectal cancer (CRC) is the third most common cancer and one of the leading cause of cancer-related death globally^1, 2^. Cancer cells expand rapidly and all solid tumors experience hypoxia due to inadequate vascularization^3^. Hypoxia plays a critical role in cancer progression via increasing angiogenesis, glycolysis, apoptotic resistance, therapy resistance, genomic instability and tumour invasion/metastasis^4, 5, 6^. Hypoxic responses are transcriptionally controlled by hypoxia-inducible factor-1α (HIF-1α), HIF-2α, and HIF-3α, which are members of the basic helix–loop–helix-PER-ARNT-SIM (bHLH–PAS) family^7, 8^. HIFs regulate multiple pathways involved in cell proliferation, survival, apoptosis, migration, metabolism, and inflammation^9, 10^. HIF-1α and HIF-2α exhibit distinct roles in colon cancers^11, 12^.

HIF-1α is positively associated with the malignant progression of various tumor entities^13, 14^. The role of HIF-1α for the pathogenesis of CRC has been studied by several groups with conflicting data. In mouse models epithelial disruption or constitutive activation of HIF-1α did not alter colon adenoma formation^15^. On the other hand, HIF-2α is essential for CRC growth and progression in cell lines and in vivo^16, 17^. The activation of intestinal epithelial HIF-2α induces a potent epithelial proinflammatory response by regulating the expression of inflammatory cytokines and chemokines^18^. While HIF-1α had no effect on tumorigenesis^19, 20, 21^, activation of HIF-2α resulted in increased tumor burden in mouse models of colon cancer^22, 23^. Our recent work has demonstrated an essential role of epithelial HIF-2α -elicited inflammation and regulation of intra-tumoral iron homeostasis as important mechanisms responsible for increase in colon cancer^24^. Also, hypoxia via HIF-2α activates YAP1 which is essential for cell growth during hypoxic stress in colon-derived cell lines^24^. Interestingly a novel, selective, potent and orally active small-molecule PT2385 was shown to selectively inhibit HIF-2α by allosterically blocking dimerization with its partner ARNT^25^. It has been shown that PT2385 is efficient to inhibit the expression of target genes in Clear Cell Renal Cell Carcinoma (ccRCC) cell lines and tumor xenografts^26, 27^. PT2385 is in Phase 2 clinical trial in patients with advanced solid tumors ccRCC. However, these strategies are not effective in long term as rapidly dividing cancer cell acquire resistance against these inhibitors very easily^28^. Therefore, there is an urgent need to increase clinical benefit of anticancer therapies by recognizing tumor cell-specific vulnerabilities. With an aim to identify hypoxic tumor-cell specific vulnerabilities we set an unbiased screen in tumor enteroids with HIF-2α overexpression. Interesting dimethyl fumarate and ferroptotic activators such as erastin, RSL3 and sorafenib were significant modulators of tumor growth and HIF-2α activation led to increase in their sensitivity. Ferroptosis is a nonapoptotic, iron-dependent form of cell death that can be activated in cancer cells by natural stimuli and synthetic agents. Ferroptosis is characterized by the loss of lipid peroxide repair capacity by the phospholipid glutathione peroxidase GPX4, the availability of redox-active iron, and oxidation of polyunsaturated fatty acid (PUFA)-containing phospholipids^29, 30^. Ferroptotic death is associated with various pathological conditions, including hepatocellular degeneration, acute kidney injury, hemochromatosis, traumatic brain injury, and neurodegeneration^31, 32^. Recently ferroptotic cell death emerged as a potent tool to target drug-tolerant persisted cancer cells^33, 34, 35^.

In the present study we utilized the combination of a hypoxic mimetic with drugs identified in our screen which led to a robust increase in the ROS, lipid ROS and decrease in glutathione production inducing cell death in comparison to their treatment alone. We identify two pathways by which HIF activated cells are highly susceptible to oxidative mediated cell death. First, activation of HIFs leads to upregulation of lipid regulatory genes in colon cancer cells as well as in mice and spots HIF-2α as a major player in inducing ferroptosis-susceptible cell state. Secondly, via a ferroptosis-independent pathway HIF-2α activation can potentiate ROS meditated cell death. These findings thus highlight HIF-2α dependent tumor cell-specific lethality and have implications for the development of novel therapeutics that could be employed for the improved treatment of colon cancer.

## RESULTS

### Drug screen identifies synthetic vulnerability to HIF2α in tumor enteroids

A small screen of known anti-cancer drugs effective in colon cancer, were assessed in in tumor enteroids^36^. Enteroids were isolated from a CDX2-CreER^T2^*Apc*^*fl/fl*^ and CDX2-CreER^T2^*Apc*^*fl/fl*^*HIF2α*^*LSL/LSL*^. Mouse. Adenomatous polyposis coli (APC) is a tumor suppressor protein mutated in more than 80% of patients with sporadic colon cancer^37^. The CDX2-CreER^T2^*Apc*^fl/fl^ mouse model enables tamoxifen inducible deletion of both *Apc* alleles in intestinal epithelial tissues. These mice were crossed with HIF2α^LSL/LSL^, which harbor an oxygen-stable HIF-2α flanked by loxP-Stop-loxP cassette (Fig.1*A*) ^38^. In CDX2-CreER^T2^-*Apc*^fl/fl^*Hif2α*^LSL/LSL^ mice tamoxifen treatment results in a robust induction of HIF-2α and disrupts *Apc* specifically in the colon epithelial cells. The enteroids isolated from this mouse and CDX2-CreER^T2^*Apc*^fl/fl^ were cultured with a panel of chemotherapeutics and growth was monitored for 5 days (Fig. 1A). Colon tumor enteroids overexpressing HIF-2α were resistant to carboplatin, cisplatin and oxaliplatin compared to enteroids from CDX2-CreER^T2^*Apc*^fl/fl^ (Fig. 1B). This data is consistent with the well-known role of HIF signaling in chemo-resistance^39^. Interestingly, erastin, RSL3, dimethyl fumarate (DMF) and sorafenib significantly reduced the growth of tumor enteroids from CDX2-CreER^T2^*Apc*^fl/fl^*Hif2α*^LSL/LSL^ in comparison to CDX2-CreER^T2^*Apc*^fl/fl^ (Fig. 1B). Erastin, RSL3 and sorafenib are classic ferroptotic activators and this data suggest HIF-2α expression may lead to high sensitivity to ferroptosis^40^.

**Figure 1.**
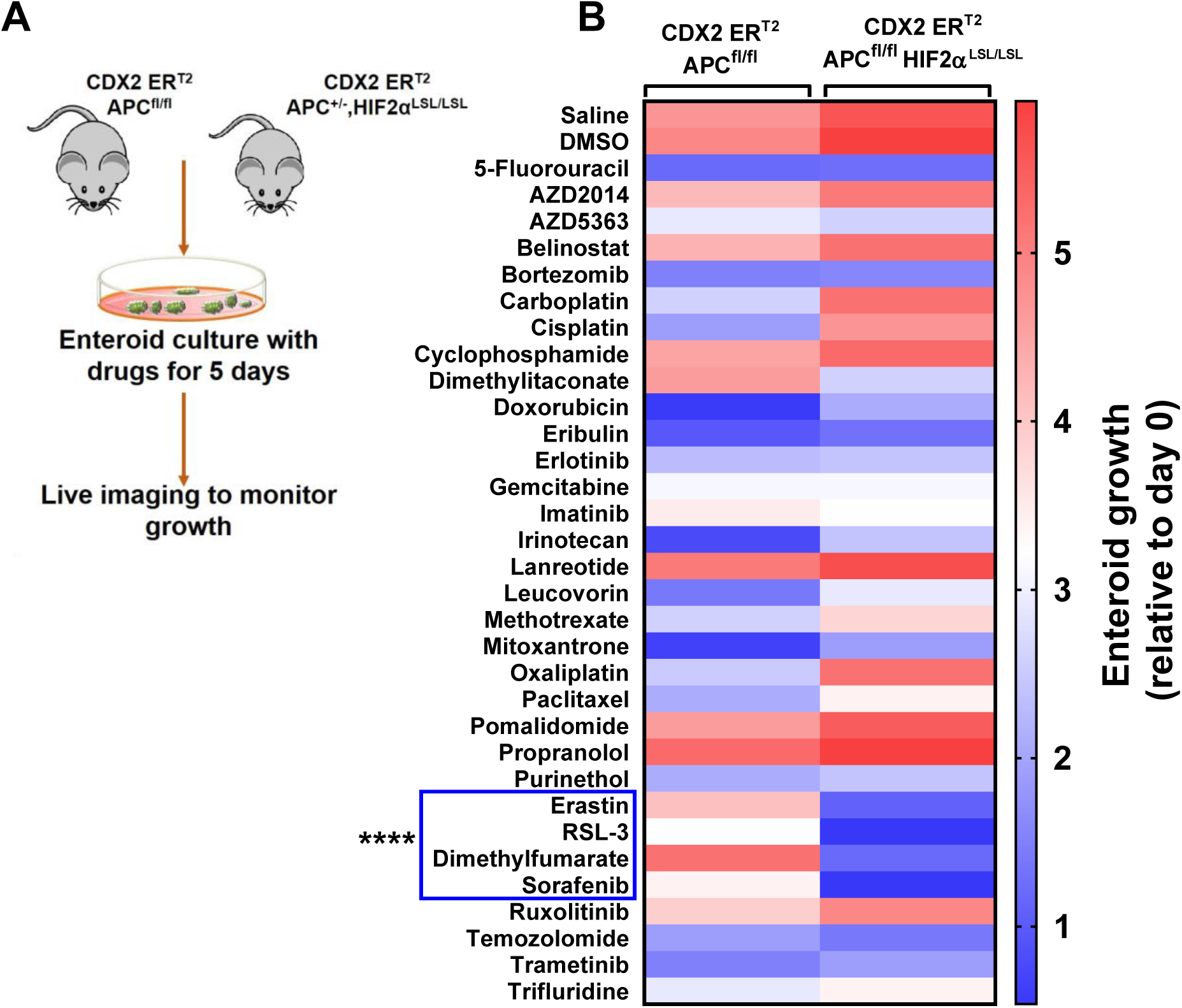
Screening of target drugs/molecules that exhibit reduction in growth of tumor enteroids. **(A)** Schematic of enteroids isolated from colon cancer HIF-2α overexpressing mouse model (CDX2ER^T2^APC^fl/fl^; HIF-2α^LSL/LSL^). Enteroids isolated from an APC^fl/fl^ and APC^fl/fl^; HIF2α^LSL/LSL^ mice were grown for 5 days with several drugs. Growth was assessed using live cell imaging (B) Heat map showing enteroid growth in the presence of various drugs. Erastin, RSL3, Dimethyl fumarate and sorafenib significantly reduced the growth of hypoxic tumors. **** P<0.0001 for the difference between CDX2ER^T2^APC^fl/fl^ compared with CDX2ER^T2^APC^fl/fl^; HIF-2α^LSL/LSL mice^

### HIF activation synergizes with ferroptotic activators in panel of colon cancer cells

Erastin and RSL3 are classical inducers of ferroptosis that were originally identified in a screening for small molecules that are selectively lethal to cancer cells^41^. Knowing that colorectal cancer cells (CRC) are solid tumors where hypoxia is a prominent feature, we first analyzed the activity of erastin and RSL3 in various human CRC cell lines, HCT116, SW480, DLD1, RKO and HT29 using MTT viability assay. Erastin (Fig 2A) and RSL3 (Fig 3A) dose-dependently induced cell death in CRC cells. To understand if activation of HIF could potentiate cell death, a hypoxia mimetic, FG4592 (Roxadustat) was assessed. FG4592 is a prolyl hydroxylase domain inhibitor which leads to stable induction of both HIF-1α and HIF-2α. Treatment of CRC cells with FG4592 for 16 hours had no effect on growth of CRC cells when compared with vehicle treated cells (Supplemental Fig. 1A). Interestingly, combination of FG4592 with ferroptotic activators-erastin (Fig. 2B) and RSL3 (Fig. 3B) potentiated cell death in CRC cells. This finding implicates the sensitivity of hypoxic colon cancer cells towards ferroptosis.

**Figure 2.**
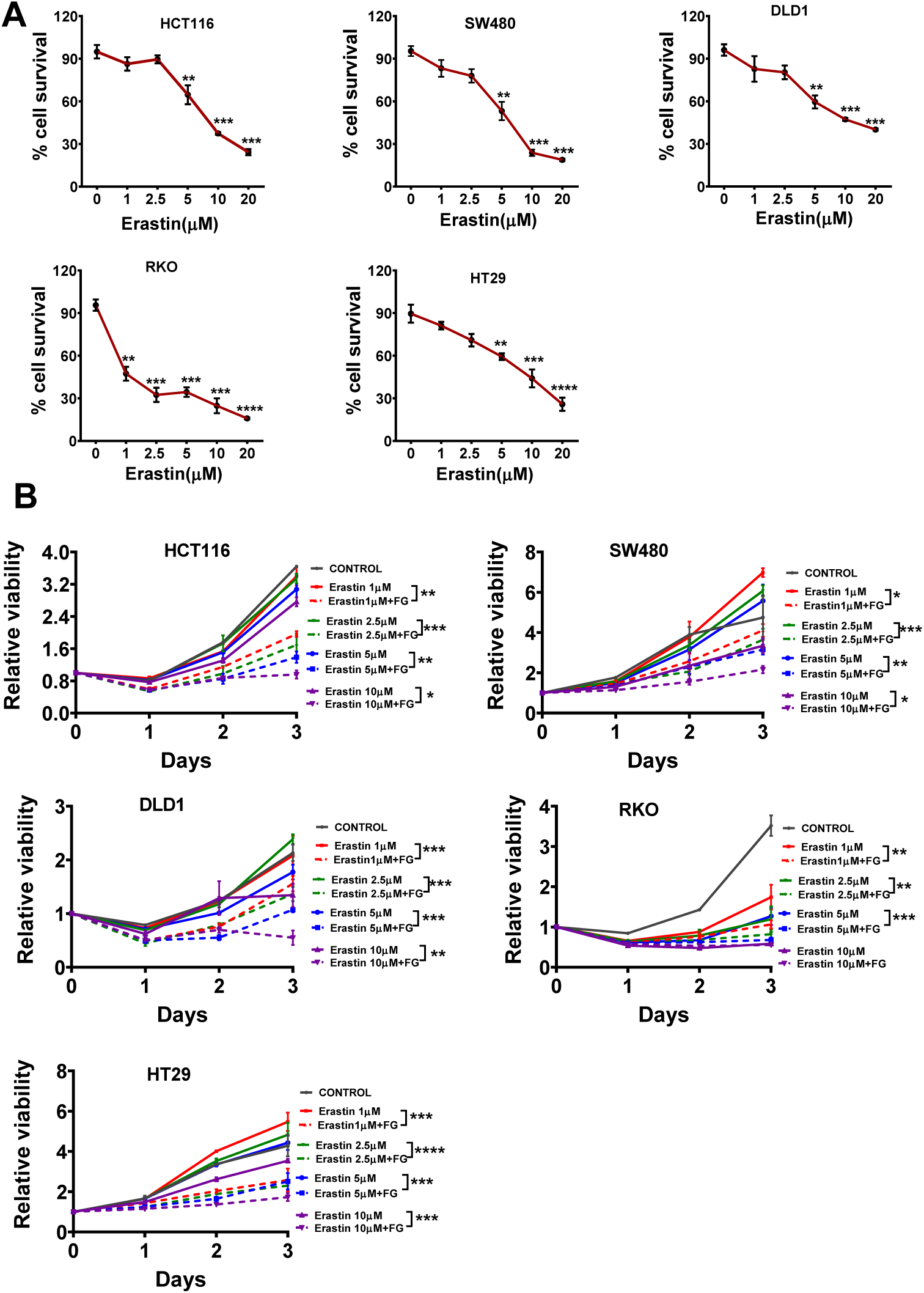
HIF-activation contributes to erastin induced cell death. HCT116, SW480, DLD1, RKO and HT29 cells were treated with 0,1, 2.5, 5 and 10 μM **(A)** erastin for 72 hours or co-treated with **(B)** FG4592 (100 μM) and varying concentrations of erastin for 3 days. Cell viabilities were assessed by the MTT assay after 72 hours or after every 24 hours in case of FG4592 and erastin treated CRC cells. Quantitative data are presented as means ± SD from three independent experiments. Statistical significance was calculated using paired-t test. *P<0.05, **P<0.01, ***P<0.001, ****p<0.0001

**Figure 3.**
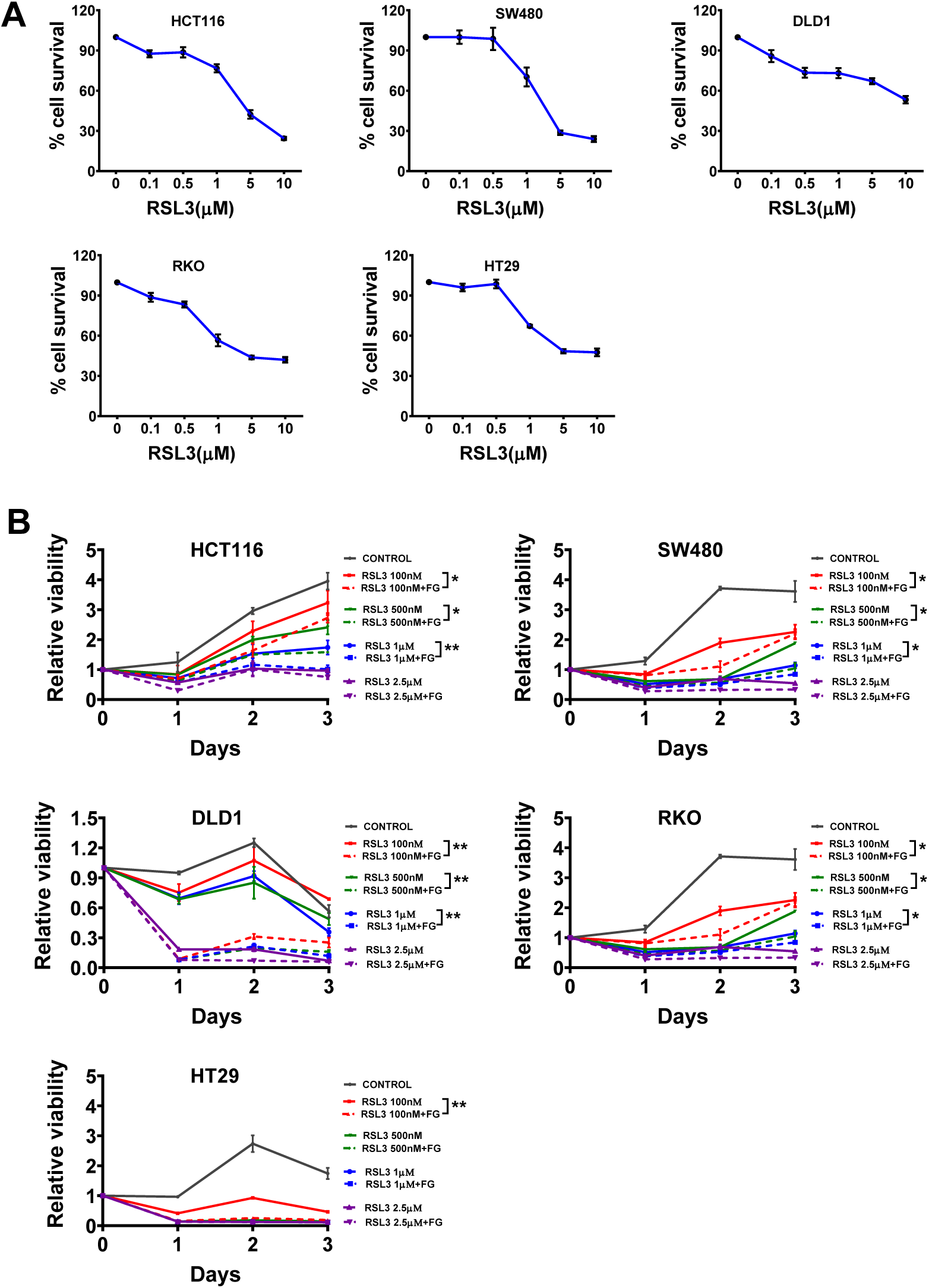
Roxadustat potentiates RSL3 induced ferroptosis. HCT116, SW480, DLD1, RKO and HT29 cells were treated with 0, 0.1, 0.5, 1, 5 and 10 μM **(A)** RSL3 for 72 hours or co-treated with **(B)** FG4592 (100 μM) with varying concentrations of erastin for 3 days. Cell viabilities were assessed by the MTT assay after 72 hours or after every 24-hour interval in case of FG4592 and erastin treated CRC cells. Quantitative data are presented as means ± SD from three independent experiments. Statistical significance was calculated using paired-t test. *P<0.05, **P<0.01, ***P<0.001, ****p<0.0001

### HIF-2α augments erastin induced ferroptosis

To understand HIF specificity (HIF-1α or HIF-2α) to ferroptosis sensitivity, HIF-1α or HIF-2α knockdown cell were utilized. The stable knockdown of HIF-1α and HIF-2α were generated in HCT116 cells and confirmed by Western analysis (Suppl. Fig. 1B and 1C). The shRNA for HIF-1α (sh_H1), HIF-2α (sh_H2) and a scrambled sequence (shScr) in HCT116 cells were treated with erastin and RSL3 either alone or in combination with FG4592 and cell survival was assessed through MTT assay. Knockdown of HIF-2α attenuated the efficacy of erastin (Fig. 4A) and RSL3 (Fig. 4B) either alone or in combination with FG4592. This data is consistent with a recent work, that suggest that HIF-2α-dependent lipid alterations increase ferroptotic cell death in renal cancer derived cell lines^31^. We analyzed the expression of two lipid genes important in ferroptosis sensitization in renal cancers, hypoxia inducible lipid droplet associated protein (HILPDA) and perilipin 2 (PLIN2). HCT116 and SW480 cells treated with FG4592 increased expression of both HILPDA and PLIN2 (Fig 4C), implicating the role of HIF-2α in driving ferroptosis through lipid accumulation. Since lipid accumulation is linked with aberrant generation of ROS, we next checked lipid ROS production through oxidation of C11-BODIPY in HCT116 and SW480 cells. Combination of FG4592 with erastin significantly increased lipid ROS (Fig. 4D) as evident by increase in fluorescence intensity of C11-BODIPY probe (Fig. 4E) which was rescued by ferroptotic inhibitor-ferrostatin-1(Fer-1) (Fig. 4D and 4E). Interestingly, FG4592 treatment alone could generate significant amount of lipid ROS in comparison to control, however FG4592 alone did not lead to cell death (Suppl. Fig. 1A). This may suggest that there is an active inhibition of ferroptosis following HIF-2α activation in cancer cells but an increase in synthetic vulnerability to erastin or RSL3. The ferroptotic target for erastin, SLC7A11 was highly increased by hypoxia and FG4592 treatment (Fig. 4F). The lipid ROS production via hypoxia may be countered by upregulation of SLC7A11 which maintains cellular redox homeostasis.

**Figure 4.**
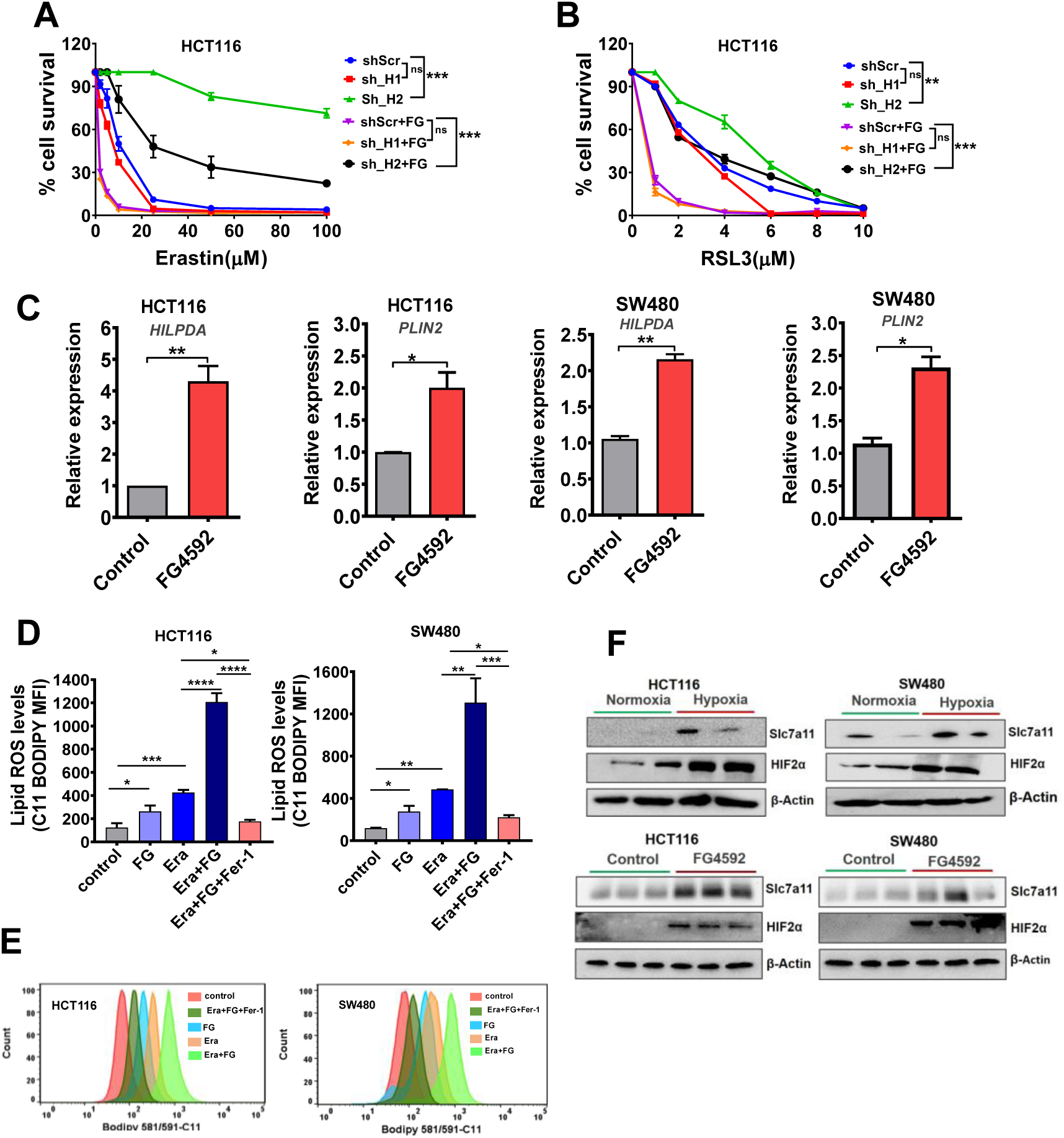
HIF2α mediates sensitivity towards erastin and RSL3 mediated ferroptosis. HIF1α and HIF2α shRNA mediated knock down HCT116 cells were treated with **(A)** 0, 10, 20, 40, 60, 80 and 100 μM erastin alone or in combination with FG4592. These cells were also treated with **(B)** 0, 1, 2, 4, 6,8 and 10 μM RSL3 and FG4592 (100 μM). Cell survival was assessed by the MTT assay after 72 hours. Quantitative data are presented as means ± SD from three independent experiments. Statistical significance was calculated using paired-t test. *P<0.05, **P<0.01, ***P<0.001, ****P<0.0001 **(C)**The mRNA levels of HILPDA and PLIN2 in HCt116 and SW480 cells treated with 100 μM FG4592 were examined by qRT-PCR. Data are means ± SEM from three independent experiments. *P<0.05, **P<0.01 **(D)** Lipid ROS levels were measured in HCT116 and SW480 cells through ferroptosis-dependent C11-BODIPY oxidation after 12-hour incubation with FG4592(100 μM), erastin (5 μM) and Ferrostatin-1 (1 μM). Data are plotted as the mean ± SD. *P* values were determined using one-way ANOVA; \**P* <0.05, \*\**P* < 0.01, ***P<0.001, ****P<0.0001 versus control cells. **(E)** A representative histogram showing the mean fluorescence intensity of C11-BODIPY 581/591 differences between the various treatments in HCT116 and SW480 ^cells **(F)** HCT116 and SW480 cells were incubated in hypoxia (1% O^2^) or were treated^ with FG4592(100μM) for HIF activation. SLC7a11 and HIF2α expression were measured using immunoblotting. Data are representative of three independent experiments.

### Temporal activation of intestinal HIF-2α with specific deletion of SLC7A11 promotes oxidative stress *in vivo*

To examine the role of HIF-2α in regulating ferroptosis *in vivo*, Villin-CreER^T2-^*Hif2α*^LSL/LSL^ were crossed with *Slc7a11*^fl/fl^. Tamoxifen treatment enables the intestinal specific deletion of SLC7A11 and overexpression of HIF-2α. (Fig. 5A). The colonic tissue from these mice were analyzed for histological changes. The deletion SLC7A11 or overexpression of HIF-2α alone where indistinguishable from control littermates. However, disruption of SLC7A11 in combination with HIF-2α overexpression led to colonic epithelial degeneration and vacuolization (Fig 5B). Lipid peroxide induced oxidative stress was measured by 4-hydroxy 2-nonenal (4-HNE) staining (Fig. 5C). The Villin-CreER^T2^*Slc7a11*^fl/fl^; *Hif2α*^LSL/LSL^ mice showed a robust increase in 4-HNE positive punctae (Fig. 5D) clearly indicating increased oxidative stress in these mice in comparison to their littermate controls. Furthermore, HILPDA mRNA levels were significantly increased in Villin-CreER^T2^-*Hif2α*^LSL/LSL^ mice compared to littermate controls (Fig. 5E). This data confirms the role of HIF-2α in ferroptosis sensitization in vivo and suggest that drugs that could induce ferroptosis could be highly efficacious in killing hypoxic cells.

**Figure 5.**
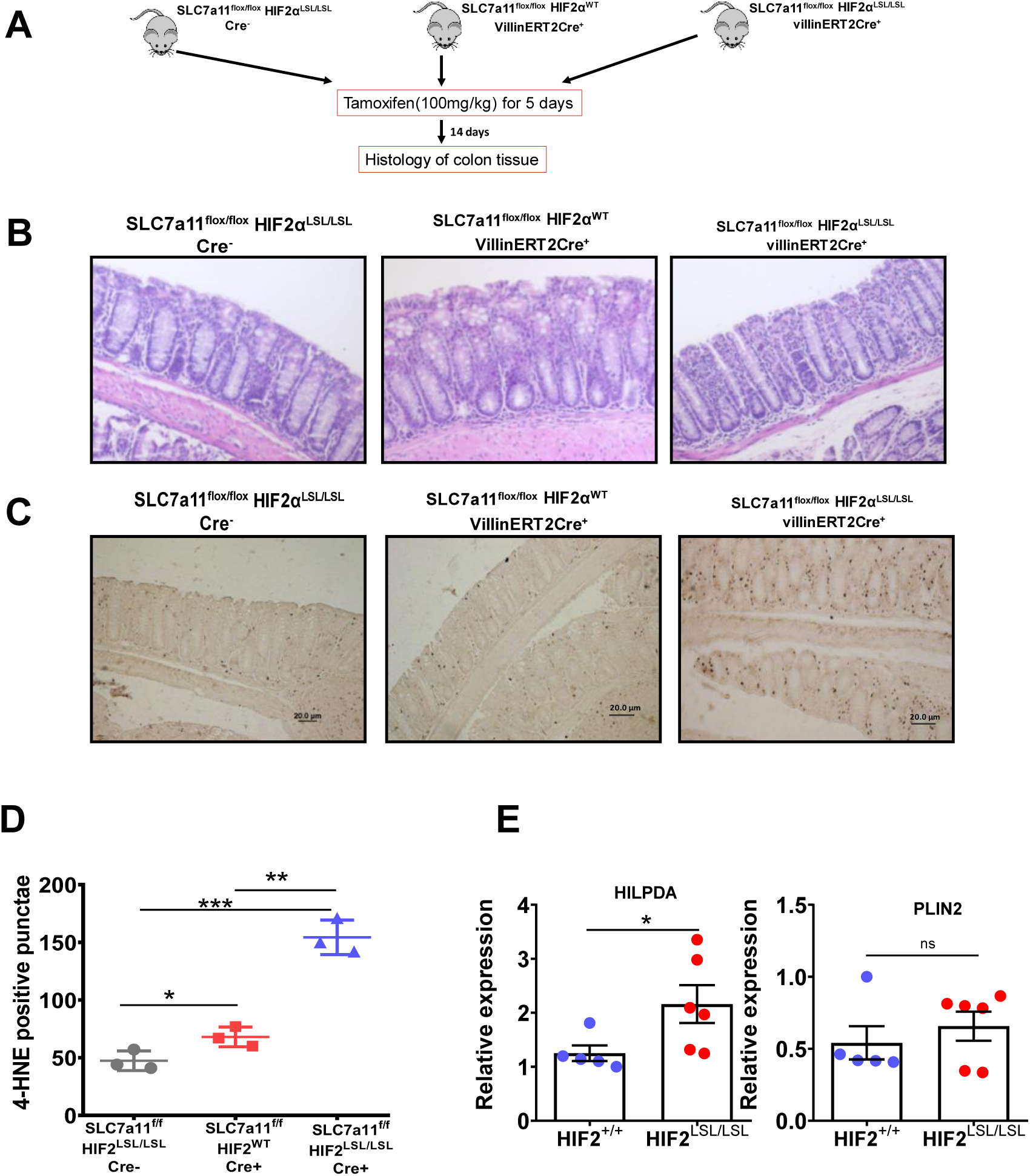
Overexpression of HIF2α and inhibition of SLC7a11 promotes oxidative stress on colonic epithelial cells in mice. **(A)** Schematic of temporal activation of intestinal HIF-2α and deletion of SLC7a11 by tamoxifen(100mg/Kg) **(B)** representative hematoxylin-and-eosin staining of colons from a SLC7a11^fl/fl^, HIF2α^lSL/LSL^ Cre-; SLC7a11^fl/fl^, HIF2α^WT^ Cre+ and SLC7a11^fl/fl^, HIF2α^lSL/LSL^ Cre+ mice. **(C)** immunohistochemistry analysis showing 4-HNE staining as a marker of oxidative stress in the 3 groups of mice as mentioned in (A). **(D)** Dot plot showing number of positively stained 4-HNE punctae or brown dots in these mice (n=3 in each group). Statistical significance was calculated using paired-t test, *P<0.05, **P<0.01, ***P<0.001 **(E)** qRT-PCR analysis for HILPDA and PLIN2 in HIF2α^WT^ (n=5) and HIF2α^LSL/LSL^ mice (n=6). Data are plotted as the mean ± SD. *P* values were determined using unpaired t-test; \**P* <0.05, ns=non-significant for the differences between compared Vil-ER^T2^ HIF2α^LSL/LSL^ compared with the littermate controls HIF2α^+/+^

### HIF activation promotes dimethyl fumarate-induced cell death in CRC cells

In the drug screen DMF was also identified as a potential molecule effective in reducing hypoxic tumor growth (Fig. 1B). DMF has previously been found to exhibit anti-tumor effects^42^. We treated a panel of colon cancer cell lines with DMF either alone or in combination with hypoxia mimetic-FG4592. Our data also showed a dose dependent inhibition of growth of colon cancer cells treated with DMF (Fig. 6A). Furthermore, FG4592 potentiates DMF-induced cell death in CRC cells (Fig. 6B). Consistent with erastin and RSL3 data, HIF-2α was essential in promoting DMF-induced cell death in CRC cells (Fig. 6 C).

**Figure 6.**
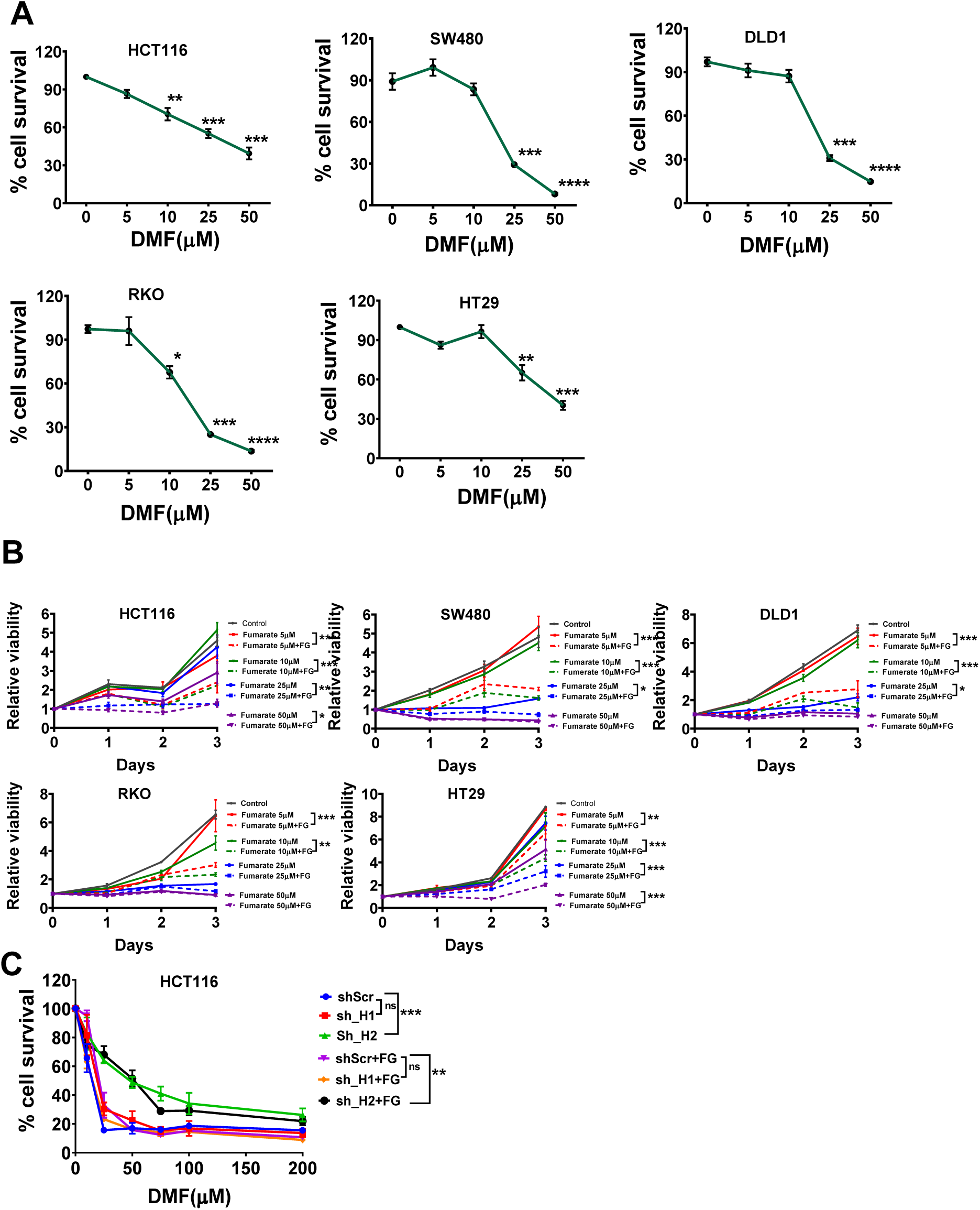
Hypoxia contributes to dimethyl fumarate (DMF) induced growth inhibition in CRC cells. HCT116, SW480, DLD1, RKO and HT29 cells were treated with 0, 5, 10, 25 and 50 μM **(A)** DMF for 72 hours or co-treated with **(B)** FG4592 (100 μM) and varying concentrations of erastin for 3 days. Cell viabilities were assessed by the MTT assay after 72 hours or after every 24 hours in case of FG4592 and DMF treated CRC cells **(C)** HIF1α and HIF2α knock-down HCT116 cells were treated with 0, 10, 25, 50, 75, 100 and 200 μM of DMF alone or in combination with FG4592 (100 μM). Cell survival was assayed by the MTT assay after 72 hours. Quantitative data are presented as means ± SD from three independent experiments. Statistical significance was calculated using paired-t test. *P<0.05, **P<0.01, ***P<0.001, ****p<0.0001

### DMF induces cell death independent of ferroptosis

Since, DMF emerged out to be effective in decreasing hypoxic tumor growth along with other ferroptotic activators (erastin, RSL3 and sorafenib) (Fig. 1B), we assessed the role of ferroptosis. HCT116 and SW480 cells treated with DMF alone or co-treated with ferroptosis inhibitors ferrostatin-1 (Fer-1) or liproxstatin-1 (Lip-1) (12,13) did not rescue the cell death induced by DMF (Fig. 7A), whereas RSL3-meidated cell death was rescued by Fer-1 and Lip-1 (Fig. 7A). Lipid ROS levels in HCT116 and SW480 cells treated with DMF or in combination with Fer-1 did not prevent the lipid ROS induction (Fig 7B) In contrast, lipid ROS indction by RSL3 was completely attenuated by Fer-1 (Fig. 7B). DMF significantly reduced cellular GSH similar to erastin and RSL3 in both HCT116 and SW480 cells (Fig. 7 C). To assess if the HIF-dependent potentiation of cell death is due to synergizing with compounds that reduce GSH, buthionine sulfoximine (BSO) a known agent to reduce GSH was assessed. The combination of DMF and BSO synergistically reduced GSH concentration in both HCT116 and SW480 cells in comparison to their treatment alone (Fig. 7D). The treatment of hypoxia mimetic also reduced cellular GSH level and synergized with BSO (Fig. 7C and 7D). However, cell viability of HCT116 and SW480 cells were not decreased with the co-treatment of BSO and FG4592 (Fig. 7E). Together, this data suggests that a novel mechanism by which hypoxia and DMF induce cell death in CRC.

**Figure 7.**
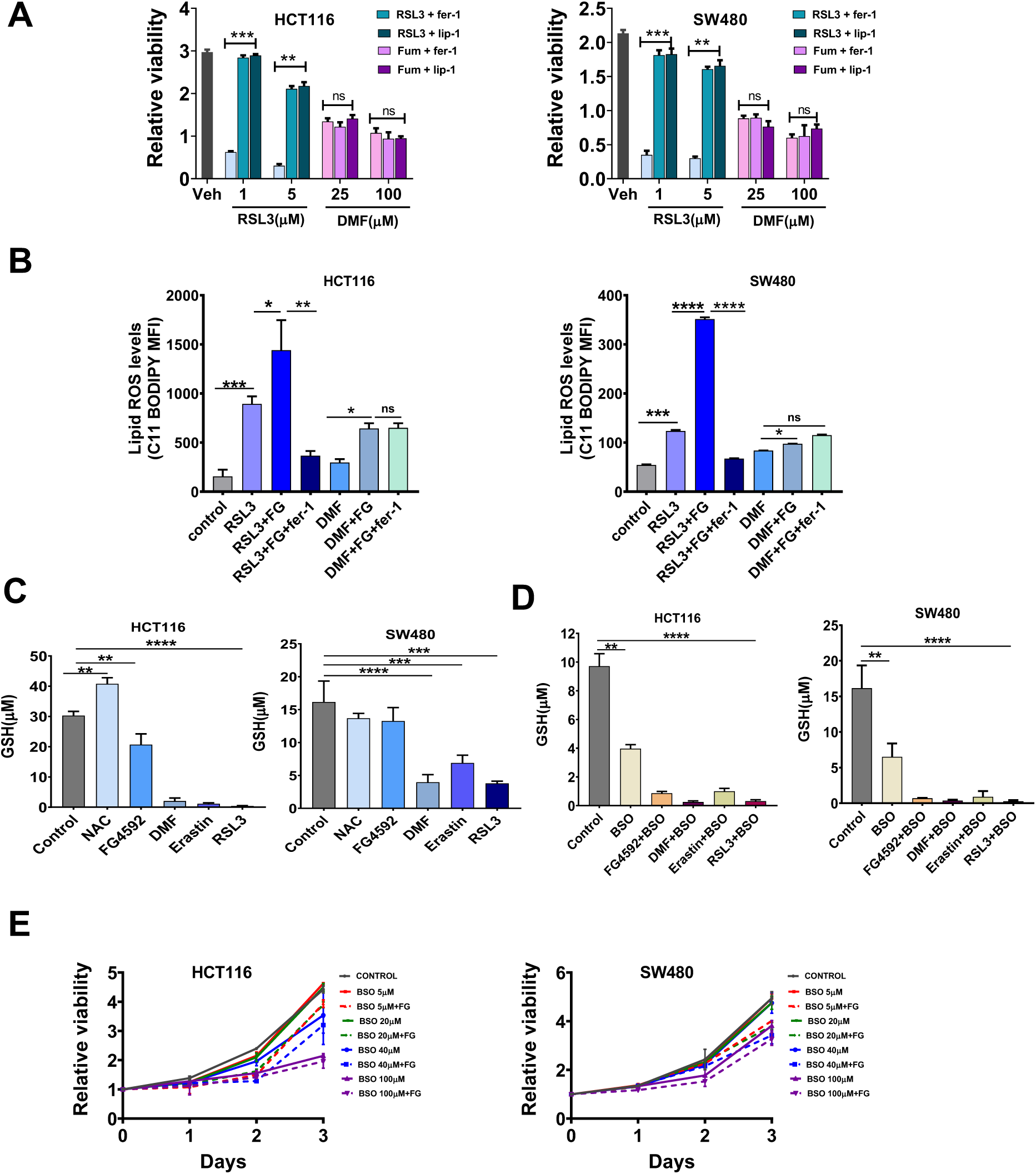
Dimethyl fumarate (DMF) is not a ferroptotic inducer for CRC cells. **(A)** HCT116 and SW480 cells were treated with RSL3 (1 and 5 μM) or DMF (25 and 100 μM) with or without ferroptotic inhibitors-ferrostatin-1 (1 μM) and Liproxstatin-1 (2 μM) for 24 hours and cell viability was assayed. Data are means ± SD from three independent experiments. Statistical significance was calculated using paired-t test. **P<0.001, ***P<0.0001, ns=non-significant. **(B)** HCT116 and SW480 cells were treated with RSL3(2 μM), DMF (50 μM) or in combination with FG4592(100 μM) with or without ferrostatin-1(1 μM) for 12 hours. Lipid ROS was determined in these cells through staining with ferroptosis-dependent C11-BODIPY581/591 dye. The dye oxidation results in green fluorescence recorded through flow cytometry. Data are plotted as the mean ± SD. *P* values were determined using one-way ANOVA; \**P* <0.05, \*\**P* < 0.01, ***P<0.001, ****P<0.0001, ns=nonsignificant. **(C)** Intracellular GSH levels in HCT116 and SW480 cells treated with N-acetyl cysteine (5 mM), FG4592(100 μM), DMF (50 μM), erastin (5 μM) and RSL3 (2.5 μM) either alone or (D) in combination with Buthionine sulfoximine (BSO) (100 μM) for 12 hours. Data are plotted as the mean ± SEM from 3 independent experiments. Statistical significance was calculated using one-way annova **P <0.01, ***P < 0.001, ****P<0.0001 versus control. **(E)** HCT116 and SW480 were treated with 0, 5, 20, 40 and 100 μM of BSO either alone or in combination with FG4592 (100 μM) for 1-3 days. Cell viabilities were assessed by the MTT assay after every 24-hour interval. Quantitative data are presented as means ± SD from three independent experiments. Statistical significance was calculated using paired-t test. ns=non-significant.

### ROS accumulation is involved in DMF-induced cell death

Our work thus far had ruled out lipid ROS and ferroptosis in increasing vulnerability of HIF-2α activated cells to DMF. To understand if the potentiation of cell death to DMF and FG4592 was mediated by oxidative stress, viability was assessed in presence of N-acetylcysteine (NAC). The viability of HCT116 and SW480 cells treated with DMF or co-treated with DMF and FG4592 was completely rescued by NAC supplementation in growth medium (Fig. 8A). Furthermore, high percentage of HCT116 and SW480 cells producing ROS was detected in FG4592 and DMF co-treatment in comparison to DMF alone (Fig. 8B). The ROS production in HCT116 and SW480 cells via DMF or DMF and FG4592 was also attenuated with NAC (Fig. 8B). This data suggests that activation of HIF leads to increased sensitivity to oxidative mediated cell death.

**Figure 8.**
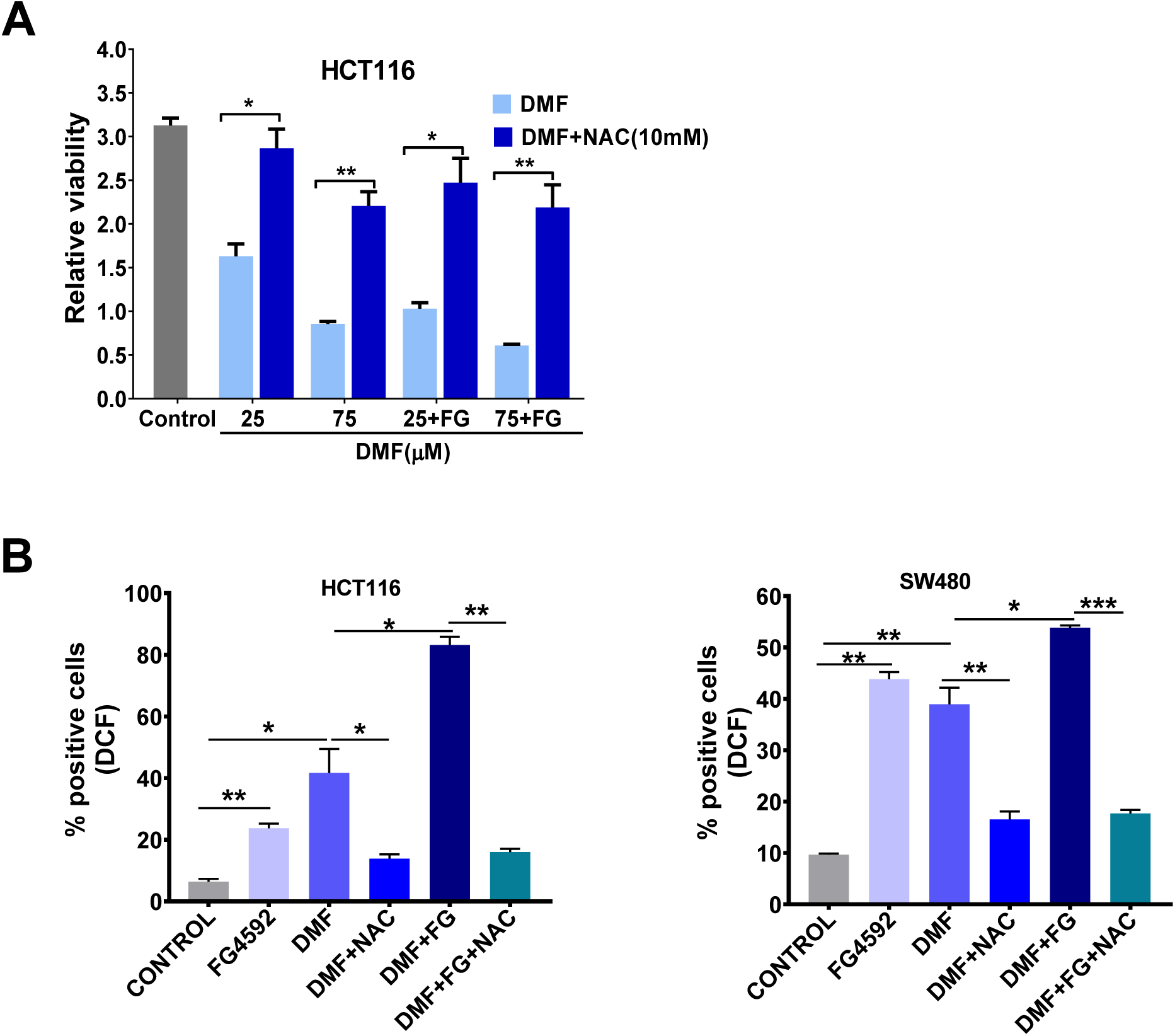
Dimethyl fumarate (DMF) induces ROS-mediated cell death in CRC cells. **(A)** HCT116 cells were treated with DMF (25 and 75 μM) or co-treated with DMF and FG4592 (100 μM) with or without N-acetyl cysteine (NAC)(5mM). The cell viability was analyzed after 72 hours with MTT assay. Data represent means ± SD from three independent experiments. Statistical significance was calculated using paired-t test. *p<0.05, **P<0.01, **(B)** HCT116 and SW480 cells were treated with FG4592(100 μM), DMF (50 μM) or DMF and FG4592 with or without NAC for 24 hours. ROS generation assay was performed using H2DCFDA and its conversion to DCF (green fluorescence) was recorded through a flow cytometer. The number of positive cells for DCF fluorescence represent the total cells with intracellular ROS. Data are plotted as the mean ± SD from 3 independent experiments. Statistical significance was calculated using one-way annova *P<0.01, **P <0.001, ***P < 0.0001.

## Discussion

Solid tumors such as CRC frequently experience inadequate supply of oxygen^43^. The cancer cells, however, have developed various adaptation strategies which enable them to grow, invade, and metastasize rapidly^44^. These mechanisms are largely employed at transcriptional level, through the stabilization of the transcription factor HIF-1α and HIF-2α^45^. Targeting hypoxic cancer cells is very difficult as the inhibitors against these transcription factors have not shown long-term efficacy. There is an urgent need to discover an intrinsic metabolic vulnerability specific to hypoxic cancer cells. Our study unravels a HIF-2α dependent synthetic lethality to colon cancer cells for erastin, RSL3 and DMF. We have previously shown that HIF-2α is a critical transcription factor involved in colon cancer progression^46^. Our drug screen data on enteroids from a hypoxic colon cancer mouse identifies erastin, RSL3, sorafenib and DMF as major molecules that could significantly reduce the growth of these hypoxic cells. Erastin and RSL3 are ferroptotic activators and have been shown to be effective in reducing growth of cancer cells including CRC cells in many of the studies. We also observe the similar effect in a panel of CRC cell lines, where either the growth inhibition or lipid ROS production by erastin or RSL3 was significantly increased by pre-treating the cells with a hypoxia mimetic. In a recent study HIF-2α has been shown to be driving vulnerability to RSL3-induced ferroptosis in clear-cell renal cell carcinoma (ccRCC)^31^. Our results also show a similar mechanism in colon cancer cells, where HIF-2α deficient CRC cells are completely resistant to high concentrations of erastin or RSL3. Moroever, we show that this vulnerability is consistent in cells and in vivo. Temporal deletion of SLC7A11 and activation of HIF-2α in intestinal epithelia cells resulted in epithelial degeneration and Lipid ROS. In the drug screen analysis, DMF was emerged out to be a potent molecule that can effectively target hypoxic tumor cell. Studies have shown that DMF mostly induced cancer cell death by GSH depletion and ROS production. Our data also show a similar role of DMF in CRC cells. Likewise, FG4592 potentiates DMF-induced ROS production and oxidative cell death, which could be restored by increase in GSH level by NAC. Our study thus unmasked the role of HIF-2α in driving synthetic lethality of hypoxic CRC cells to inhibitors that can produce oxidative stress. This intrinsic vulnerability could be efficiently combined with anti-cancer therapies to target highly resistant hypoxic tumors.

## Methods

### Animal Experiments

All animal studies were carried out in accordance with Institute of Laboratory Animal Resources guidelines and approved by the University Committee on the Use and Care of Animals at the University of Michigan (IACUC protocol number: PRO00008292) For all experiments, male and female mice, 6 to 8-weeks of age were used. All mice are a C57BL/6 background maintained in standard cages in a light- and temperature-controlled room, and were allowed a standard chow diet and water ad libitum. Villin-CreER^T2^ *Hif2α*^LSL/LSL^ and *Apc*^fl/fl^ mice have been previously described^24, 38^. These mice were crossed with the colon specific Cre to generate the CDX2-CreER^T2^-*Apc*^fl/fl^; *Hif2α*^LSL/LSL^ mice. Correctly targeted ES cells in which exon 3 of *Slc7a11* was flanked by Lox-P (*Slc7a11*^fl/fl^) sites were generated by the International Mouse Phenotyping Consortium. *Slc7a11*^fl/fl^ mice were also crossed to the Villin-CreER^T2^ mice and further crossed to the *Hif2α*^LSL/LSL^ mice *to generate the* Villin-CreER^T2^*-Slc7a11* ^fl/fl^;*Hif2α*^LSL/LSL^ mice. For all experiments littermate controls were used and the Cre was actvated by I.P injection with tamoxifen in corn oil (100 mg/kg) for three consecutive days and were euthanized 1 week or 2 week following the last tamoxifen treatments.

### Cell lines and reagents

HCT116, SW480, DLD1, RKO and HT29 cells were obtained from ATCC and grown in DMEM with L-glutamine, D-glucose and sodium pyruvate (GIBCO) supplemented with 10% heat-inactivated FBS and 1% antibiotic-antimycotic mix (Invitrogen). All cells were maintained in a humidified environment at 37 °C and 5% CO2 in a tissue culture incubator. DMF and BSO was purchased from Sigma-Aldrich (St. Louis, MO, USA). Erastin, RSL3, FG4592, Ferrostatin-1(Fer-1), Liproxstatin-1(Lip-1) and N-acetyl cysteine (NAC) were purchased from Cayman chemical.

### C11-BODIPY lipid ROS measurement

1×10^6^ HCT116 or SW480 cells were seeded in 12-well plates and allowed to adhere overnight at 37 °C. The day before the experiment, cells were treated with DMSO(vehicle), erastin (5 μM), RSL3 (2 μM) or DMF (50 μM), ferrostatin −1(Fer-1) (1 μM) with or without FG4592 (100 μM) and incubated for 12 h at 37 °C. Cells were harvested using PBS-EDTA(5mM), buffer, washed once with HBSS and suspended in HBSS containing 5 μM C11-BODIPY (ThermoFisher) and incubated at 37 °C for 30 min. Cells were pelleted, washed and resuspended in HBSS. Fluorescence intensity was measured on the FITC channel using Beckman Coulter MoFlo Astrios. A minimum of 20,000 cells was analyzed per condition. Data was analyzed using using FlowJo software (Tree Star, Ashland, OR, USA). Values are expressed as mean fluorescence intensity (MFI).

### ROS detection assay

The cell-permeable free radical sensor carboxy-H2DCFDA (Invitrogen) was used to measure intracellular ROS levels. Cells treated with DMF (50 μM), FG4592(100 μM) with or without N-acetyl cysteine (NAC) were harvested by ice-cold PBS-EDTA(5mM) buffer and incubated with 10 μM carboxy-H2DCFDA in PBS at 37°C for 30-45 min. The cells were washed, resuspended in PBS and analyzed using Beckman Coulter MoFlo Astrios flow cytometer. Data were analyzed using FlowJo software. Values are expressed as the percentage of cells positive for DCF fluorescence.

### MTT assay

2000-3000 cells were seeded in a 96-well plate and allowed to adhere overnight at 37 °C. The next day cells were treated with FG4592(100 μM) for 16 hours in co-treatment conditions. The cells were then treated with different agents as indicated in figures. 24-hours following treatment a Day 0 reading was taken. Following the Day 0 read, the corresponding treatment and readings were taken every 24-hours for 72-hour assay. Cells were incubated for 45 minutes with Thiazolyl Blue Tetrazolium Bromide (Sigma) then solubilized with dimethyl sulfoxide. Absorbance was taken at 570nm. All reads were taken in technical triplicates.

### GSH assay

HCT116 and SW480 cells plated in a 96-well plate were treated with different agents or the vehicle for 24 hours. The GSH concentrations were determined using the GSH-Glo Glutathione Assay Kit (Promega) as per the manufacturer’s instructions. The luminescence-based assay is based on the conversion of a luciferin derivative into luciferin in the presence of GSH, catalyzed by glutathione *S*-transferase. Luciferase expression was then measured on a Synergy Mx Microplate Reader (BioTek). The signal generated in a coupled reaction with firefly luciferase is proportional to the amount of GSH present in the sample. The concentration was determined through a standard curve using GSH standard solution provided with the kit.

### Western blotting

HCT116 and SW480 cells were seeded at 1m/mL density in a 6 well plate in triplicates for each condition and allowed to adhere overnight. Cells were then treated with FG4592 (100 μM) or incubated in hypoxic chamber (1% O_2_) for 16 hours. Cells were lysed with radioimmunoprecipitation (RIPA) assay buffer with added protease (1:100 dilution; Sigma) and phosphatase (1:100 dilution; ThermoFisher Scientific) inhibitors. After cell lysis, solubilized proteins were resolved on 10% SDS-polyacrylamide gels and transferred to nitrocellulose membrane, blocked with 5% milk in TBST and were immunoblotted with the indicated primary antibodies made at 1:1000 dilution in blocking solution for HIF-1α(Abcam), HIF-2α (Bethyl lab), SLC7A11(Abcam) and β-actin (Proteintech).

### Histology and 4HNE staining

Colonic tissues were rolled and fixed with PBS-buffered formalin for 24-hours followed by embedding in paraffin. 5μM sections were stained for Hematoxylin-and-eosin(H & E) and mounted with permount mounting medium(Fisher scientific). For, 4 hydroxy-2-noneal/4HNE staining, paraffin-embedded tissue sections were subjected to antigen retrieval, followed by blocking with 5% goat serum in PBS and probed with primary antibody against 4HNE (1:200 dilutions, BS6313R, Bioss). Sections were then washed three times with PBST and were incubated with HRP conjugated anti-rabbit IgG (1:500 Dilution, Cell Signaling technology) for 1 h. Sections were then washed three times with PBST, and incubated with DAB substrate solution to sufficiently cover them. After the sample color turned brown, the reaction was stopped by distilled water and dehydration steps were carried. The slides were mounted using permount mounting medium.

### Enteroid culture with drugs

Mouse colonic crypts were isolated using a previously described method^47^. Colon was isolated from CDX2-ER^T2^CreApc^fl/fl^ HIF2α^LSL/LSL^ and CDX2-ER^T2^CreApc^fl/fl^ mice and cut in to 1cm pieces. Tissue was incubated in 10mM DTT for 15 minutes at room temperature. Tissues were rinsed with DPBS supplemented with gentamicin and primocin. Tissue was incubated with slow rotation at 4C for 75minutes in 8mM EDTA. EDTA was removed and tissue was put through three cycles of snap-shakes to release crypts. Isolated crypts were spun down and collected in cold LWRN medium. Crypts were then plated in matrigel (Corning) in 96-well culture plates in LWRN media and imaged using Image express Micro. Enteroids were treated with indicated drugs/inhibitors (10μM) and growth was monitored after 5 days.

### RNA isolation and qPCR analysis

HCT116 and SW480 cells were seeded at 1m/mL density in a 6 well plate in triplicates for each condition and allowed to adhere overnight. Cells were then treated with FG4592 (100 μM) for 16 hours. RNA was isolated from cultured tissue or using TRIzol chloroform extraction method. RNA was reverse transcribed using MMLV reverse transcriptase (ThermoFisher). qPCR analysis was done using indicated primers and Radiant Green qPCR master mix (Alkali Scientific Inc.)

### Statistical Analysis

Data represent the mean ± s.e.m or s.d. in case of viability experiments. Data are from three independent experiments measured in triplicate, unless otherwise stated in the figure legend. For statistical analyses, Student’s *t*-tests were conducted to assess the differences between two groups. One-way ANNOVA was used for multiple treatment conditions. A *P* value of less than 0.05 was considered to be statistically significant. All statistical tests were carried out using Prism 8 software (GraphPad)

