## Supplemental Figures for "Hypoxia inducible factor-2α increases sensitivity of colon cancer cells towards oxidative cell death"

### Suppl. Fig. 1

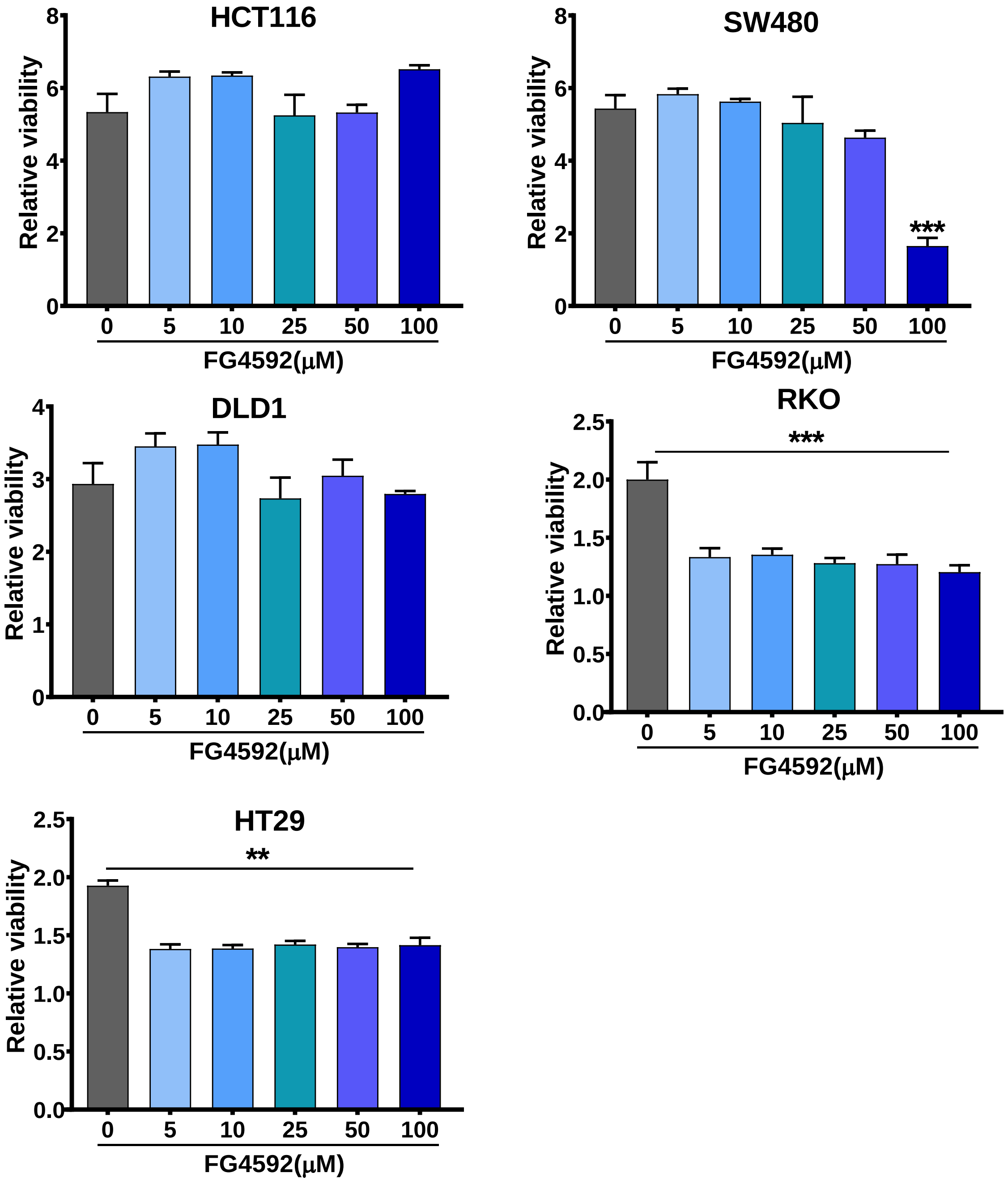


**Suppl. Fig. 1 Effect of FG4592 on growth of CRC cells.** HCT116, SW480, DLD1, RKO and HT29 cells were treated with 0,5,10, 25, 50 and 100 μM for 72 hours. Cell viability was assessed by the MTT assay after 72 hours. Quantitative data are presented as means ± SD from three independent experiments. Statistical significance was calculated using paired-t test. **P<0.01, ***P<0.001

### Suppl. Fig. 2

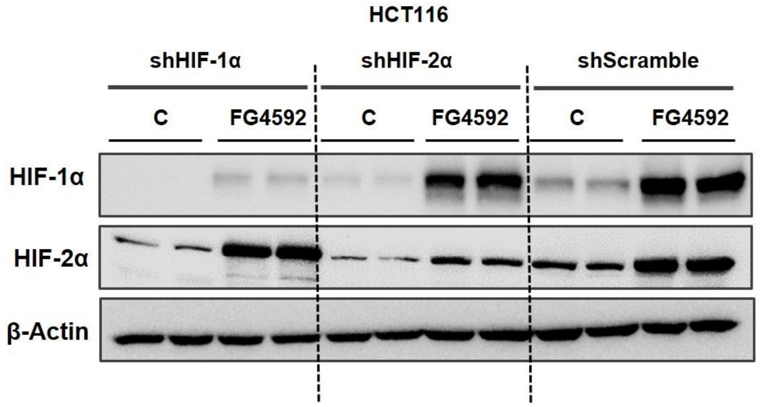


**Suppl. Fig. 2 Protein expression in shRNA mediated knock-down cells.** sh HIF-1α, sh HIF-2α and non-target sh scramble HCT116 were treated with FG4592(100μM) for HIF activation. HIF-1α and HIF2α expression was measured using western blotting. Data are representative of three independent experiments
